# Sex-Specific Metabolic Programming in Human Neutrophil Subsets

**DOI:** 10.1101/2025.08.07.669054

**Authors:** Anjali S Yennemadi, Faye K Murphy, Joseph Keane, Gina Leisching

**Author notes:** Corresponding author: Gina Leisching.

## Abstract

**Background:** Sex differences in immune responses are well-documented, with females exhibiting more robust immunity against infections but higher susceptibility to autoimmune diseases, while males often demonstrate more severe inflammatory pathology. Neutrophils which are key players in the innate immune response, display sex-based functional differences, but whether these extend to metabolic programming, particularly in low-density neutrophils (LDNs), remains unknown.

**Methods:** We isolated LDNs and normal-density neutrophils (NDNs) from healthy human donors using density gradient centrifugation and negative selection. Cellular metabolism was assessed via Seahorse XF analysis (measuring oxygen consumption rate, OCR, and extracellular acidification rate, ECAR), alongside flow cytometry for maturity markers (CD16hi/lo).

**Results:** Male LDNs exhibited significantly higher basal OCR and ATP production than female LDNs, while no sex differences were observed in NDNs. Strikingly, male LDNs had higher OCR and glycolysis than their matched NDNs, whereas female NDNs were more oxidative than their LDNs. These metabolic differences were independent of neutrophil maturity, as CD16hi frequencies did not differ between subsets or sexes.

**Conclusions:** Our study reveals, for the first time, subset-specific sexual dimorphism in neutrophil metabolism whereby male LDNs adopt a hypermetabolic (oxidative/glycolytic) phenotype, while female NDNs retain higher oxidative capacity. This reprogramming occurs independently of developmental stage, suggesting sex hormones or epigenetic regulation may drive these differences. These findings provide a metabolic basis for sex-biased immune responses and highlight the need for sex-stratified approaches in neutrophil-targeted therapies.

**Plain English Summary:** Men and women fight infections differently, and our study could reveal why—their immune cells produce energy in distinct ways. We examined two types of neutrophils (infection-fighting cells), normal cells (NDNs) and low-density cells (LDNs) found in inflammation. Using advanced metabolic measurements, we discovered that men’s LDNs use more oxygen and generate more energy than women’s. Surprisingly, while men’s LDNs were more active than their normal neutrophils, women showed the opposite—their normal neutrophils were more energetic than their LDNs. These differences weren’t due to cell maturity, suggesting biological distinctions between sexes. This may help explain why men often have worse outcomes in diseases like sepsis (where oxygen-driven inflammation dominates), while women are more prone to autoimmune diseases like lupus. Our findings could lead to better sex-specific treatments by dampening overactive immune responses in men or adjusting metabolism in women to prevent autoimmune flares. This research highlights why medical studies must consider sex differences, as one-size-fits-all treatments may miss key biological variations.

**Highlights:**

- CD16hi frequencies show no sex differences, however female (but not male) NDNs contain significantly more mature CD15+ CD16hi cells than their LDNs, revealing female-specific maturation differences between subsets.
- First report of sex-specific metabolic differences in LDNs where males exhibit significantly higher basal respiration and ATP production than female LDNs.
- Male LDNs show higher OCR than matched NDNs, while female NDNs are more oxidative than their LDNs.

## Introduction

Sex differences in immune function and metabolism are well-documented, with females exhibiting a higher prevalence of autoimmune diseases, while males often demonstrate increased susceptibility to severe infections [1-3][4-7]. These differences arise from a complex interplay of factors, including sex chromosomes, hormonal regulation, epigenetics, and environmental influences [8]. Androgens and progesterone primarily act as an immunosuppressors, with androgens taking on a protective role in the immune response, whereas estrogen can enhance the immune response[9, 10]. For instance, sex hormone-binding globulin modulates estrogen signaling in lymphocytes and has been implicated in metabolic disease pathogenesis[11]. Beyond hormones, emerging evidence highlights additional contributors to immune sexual dimorphism, such as the transcription factor VGLL3, which regulates autoimmune-associated gene expression [12].

Low-density neutrophils (LDNs), a subset co-isolating with peripheral blood mononuclear cells (PBMCs) in density gradients, have garnered attention for their roles in chronic disease[13], pregnancy[14], and even healthy states [15]. LDNs exhibit a density of ∼1.077 g/mL, distinct from normal-density neutrophils (NDNs, ∼1.083 g/mL) [19], and are hypothesized to represent an immature granulocyte population (e.g, promyelocytes/myelocytes). Their heterogeneity is increasingly recognized, with mature and immature phenotypes potentially coexisting in a context-dependent manner [15]. While LDNs correlate with disease severity in some settings such as SLE[16], they may also exert a protective role[17]. Notably, murine studies report that immature LDNs possess elevated bioenergetic capacity and a heightened propensity for NETosis compared to NDNs [18-20], but whether similar metabolic distinctions exist in humans—and whether they differ by sex— remains unexplored.

Current research has shown that male neutrophils are less active, less prone to LPS-stimulated NETs, but are more responsive to drug treatments[21-23]. Young adult total neutrophils have also shown have increased oxidative phosphorylation (OXPHOS) with these differences being attributed to a lack of estradiol. Despite these advances, it is still unknown whether sub-set specific metabolic differences occur. This is noteworthy considering the established link between cellular metabolism and immunity. Our aim was therefore to determine whether bioenergetic differences existed within each neutrophil subtype in males versus females.

## Materials and Methods

### Ethical Approval and Blood Collection

Healthy blood was collected from 22 consented individuals (n=11 males, n=11 females) and approved by the St James’s Hospital/Tallaght University Hospital Joint Research Ethics Committee. Healthy donors were defined as age and sexed matched. Blood was collected in Lithium-Heparin coated collection tubes and processed within the first hour.

### LDN/NDN isolation from whole blood

Total neutrophils were isolated from 12 mL whole blood from healthy donors using the EasySep™ Direct Human Neutrophil Isolation Kit (StemCell Technologies #19666) via negative selection. Low-density neutrophils and normal-density neutrophils were further separated by discontinuous Percoll density gradient centrifugation as we have previously described[24]. Isolated cells were resuspended in EasySep™ Buffer (StemCell Technologies #20144), counted, and processed further for downstream assays. Purity was assessed through flow cytometry and were found to be 93% pure.

### Agilent Mitostress Test

Cells were plated into a Seahorse XFp cell culture microplate (Agilent Technologies, Inc.) at a seeding density of 1.25 × 10^5^ cells/well, immediately after counting. Mitochondrial function was assessed using the Cell MitoStress Test (Agilent Technologies, Inc.) on the day of isolation. All values were normalized using the Crystal Violet dye extraction growth assay and the Agilent Wave Desktop 2.6 Software (https://www.agilent.com) was used for analysis. LDNs and NDNs were sequentially treated with oligomycin (1.5µM), carbonyl cyanide-4-(trifluoromethoxy) phenylhydrazone (FCCP; 0.5µM) and rotenone + antimycin A (Rot/AA; 0.5µM).

### Flow Cytometry

Isolated LDNs and NDNs were resuspended in phosphate-buffered saline (PBS) at a concentration of 1 × 10^6^ cells/ml for subsequent immunophenotyping by flow cytometry. These were surface stained with an antibody cocktail containing CD14, CD86, CD15, CD16 antibodies, Fc block, and viability stain for 15 minutes in the dark, at room temperature (RT). The cells were then fixed using 2% paraformaldehyde (PFA) for 15 minutes in the dark at RT. Finally, the isolated LDNs and NDNs were resuspended in PBS to be acquired on a FACS Canto II (BD Biosciences). Data analysis was performed using FlowJo software (Version 10.10.0, FloJo LIC, Oregon, USA).

### Statistical analysis

Statistical analyses were performed using GraphPad Prism 10 software. Statistically significant differences between two normally distributed groups were determined using Student’s paired t-tests with two-tailed P-values. Differences between three or more groups were determined by one-way ANOVA with Tukey’s multiple comparisons tests. P-values of < 0.05 were considered statistically significant.

## Results

### Female LDNs are more immature than their NDN counterparts

We observed no significant difference in the absolute number of LDNs between males and females (Fig. 1A). No sex difference was seen in the CD16hi frequency of CD15+ cells in LDNs or NDNs (Figure 1D). Female NDNs had a significantly higher frequency of CD16hi CD15+ cells compared to their LDNs counterparts. This difference was not seen in males (Figure 1D). Additionally, there was significant difference in the CD16lo/int frequency of CD15+ cells between female LDNs and NDNs, which was also not observed in males (Fig. 1E). Additionally, no CD16lo/int differences were seen when compared between the sexes (Fig. 1E).

**Figure 1.**
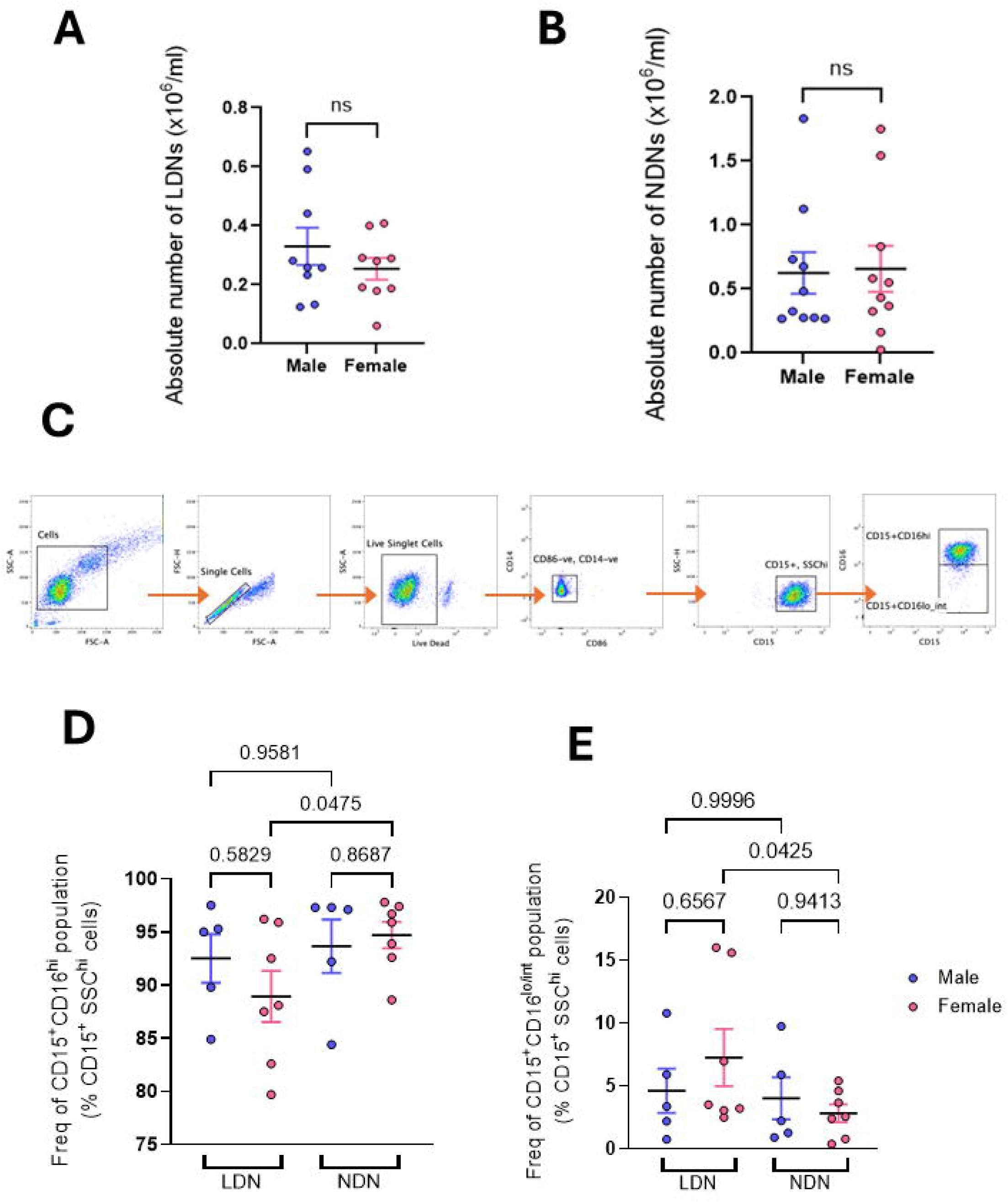
Female LDNs have fewer CD16hi CD15+ cells and more CD16lo/int compared to their NDN counterparts, an observation not observed in males. **A**. Absolute numbers of LDNs in males and females remain stable. **B**. Absolute numbers of NDNs in males and females, show no significant difference. **C**. Representative flow cytometry gating strategy for identifying CD16 levels in neutrophils. **D**. CD16hi frequency of CD15+ in male and female LDNs and NDNs, CD16hi levels are significantly higher in female NDNs compared to female LDNs. **E**. CD16lo/int frequency of CD15+ cells in LDNs and NDNs from both males and females, CD16lo/int levels are higher in female LDNs in comparison to female NDNs, a difference not seen in males. Mann-Whitney t-test or two-way ANOVA with multiple comparisons, n= 5-10.

### LDNs from males exhibit increased oxygen consumption rate

Using the Mitostress test, we wanted to determine the bioenergetic capacity of LDNs and NDNs of males vs. females (Fig. 2 and 3). We observed that male LDNs exhibited a significantly higher basal oxygen consumption rate (OCR) compared to female LDNs, indicating greater mitochondrial oxidative activity (Figure 1A and C). In contrast, basal ECAR (a surrogate for glycolysis) did not differ between sexes (Figure 1B and C). Further analysis revealed that male LDNs had elevated ATP-linked respiration, while maximal respiration remained unchanged (Figure 1C). Energetic phenogram profiling demonstrated an overall more energetic profile in male LDNs whereas female LDNs trended toward a more glycolytic phenotype (Figure 1D). These findings suggest intrinsic sex-based metabolic programming in LDNs, independent of cell frequency.

**Figure 2.**
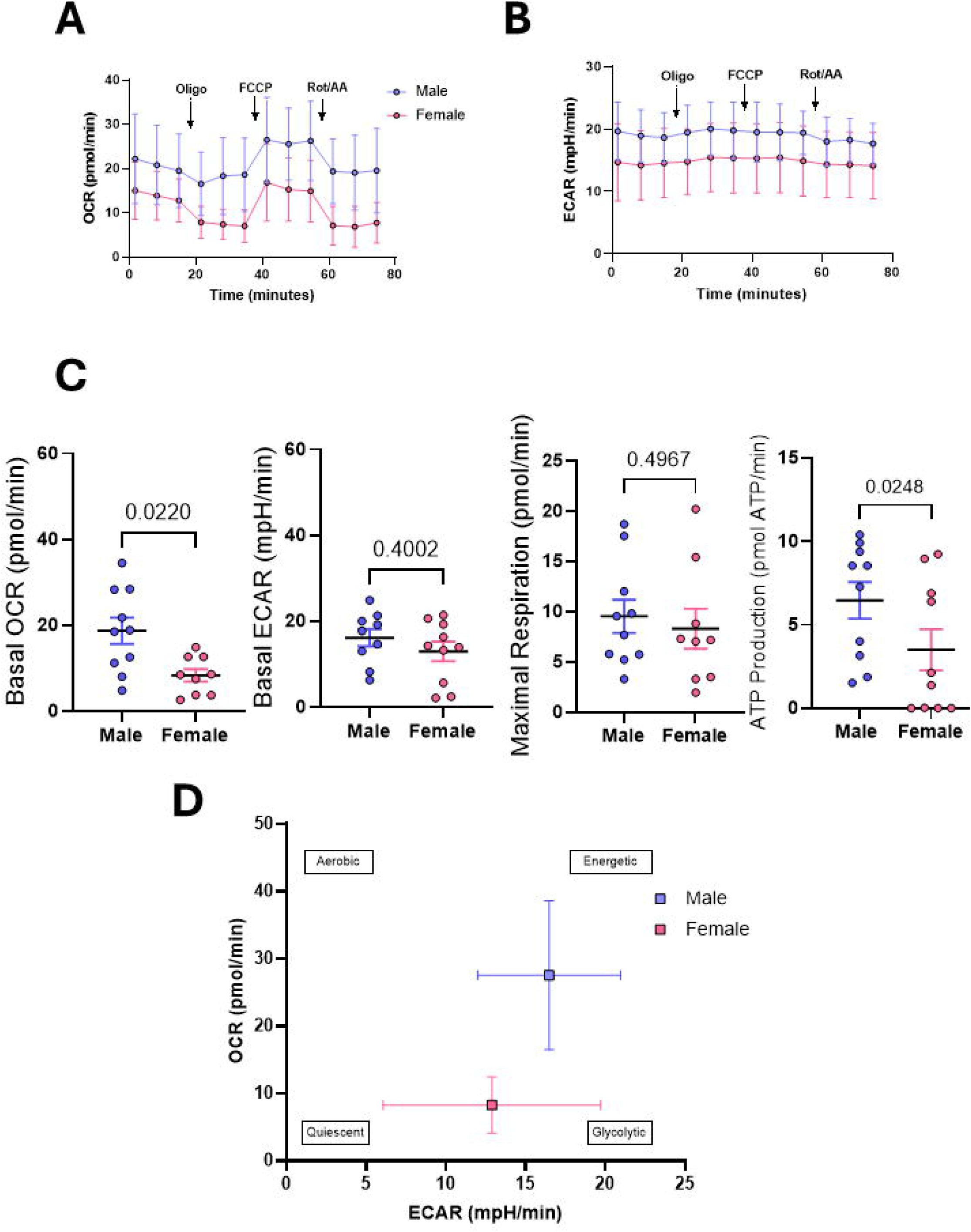
LDNs from males exhibit increased basal OCR, ATP production and overall bioenergetic capacity compared to female LDNs. **A**. LDNS were isolated assessed using the Agilent Mitostress test to assess mitochondrial fitness and oxygen consumption rate. **B**. ECAR was assessed simultaneously in LDNs from males and females. **C**. Metabolic parameters such as basal OCR and ATP production were significantly higher in male vs. Female LDNS. No changes in basal ECAR and maximal respiration was observed. **D**. Energetic phenogram depicting male LDNs as having a more energetic phenotype compared to female LDNs which have a more glycolytic phenotype in comparison. Mann-Whitney T-test, n=9-11.

**Figure 3.**
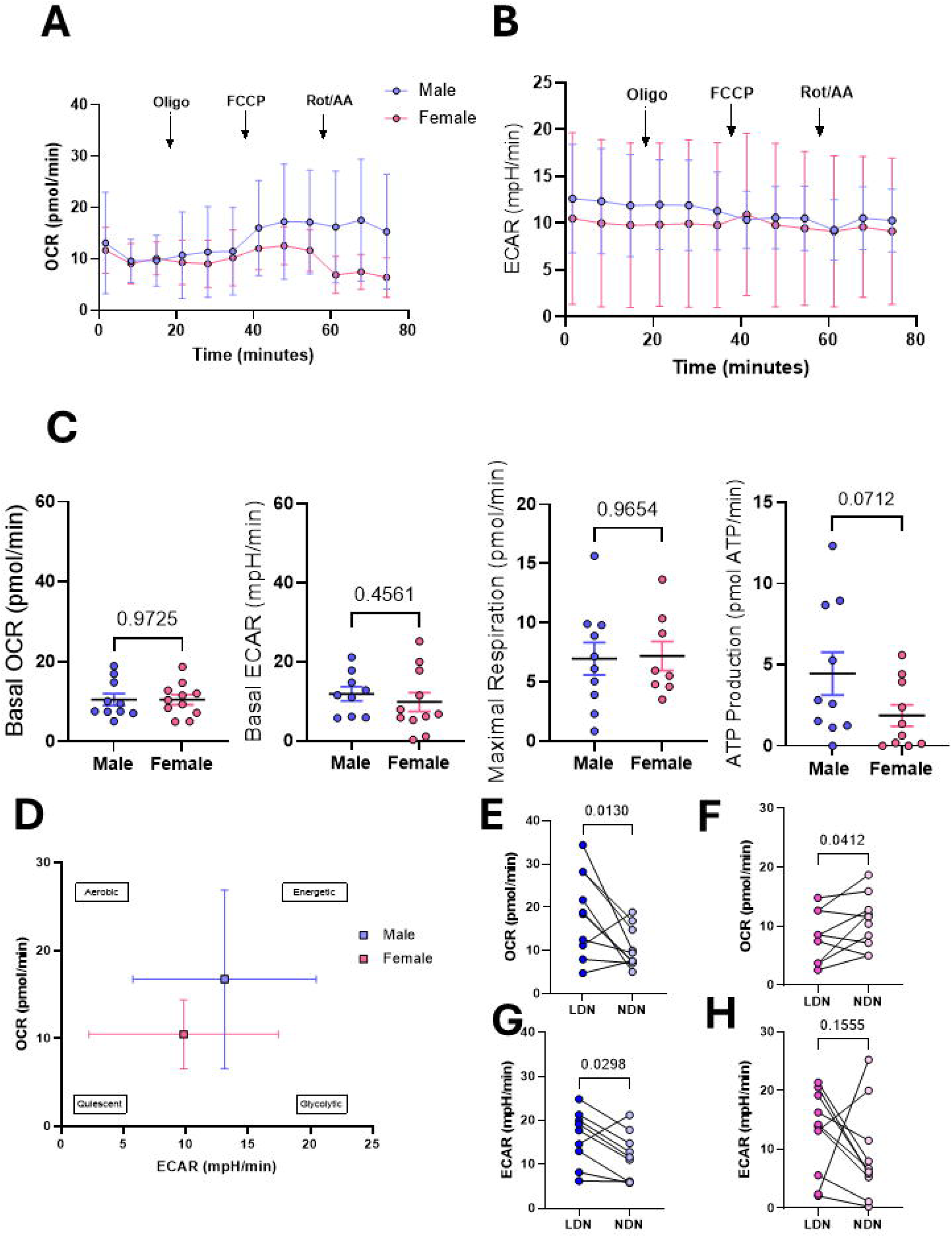
NDN bioenergetics to not differ between males and females. **A**. NDNS were isolated assessed using the Agilent Mitostress test to assess mitochondrial fitness and oxygen consumption rate. **B**. ECAR was assessed simultaneously in NDNs from males and females. **C**. No changes in basal OCR, basal ECAR, maximal respiration and ATP production were observed in male vs. Female NDNS. **D**. Energetic phenogram depicting male NDNs as having a more energetic phenotype compared to female NDNs which have a more quiescent phenotype in comparison. **E**. Male LDNs have a higher OCR compared to their NDN counterparts, which was observed to be the reverse for **F**. female LDN OCR. **G**. Male LDNs had a significantly higher basal ECAR compared to their NDN counterparts, a trend which was not observed in **H**. female LDN/NDNs. Mann-Whitney T-test or paired t-test, n=9-11.

### Normal-Density Neutrophils do not exhibit metabolic Sex Differences

No differences were detected in basal OCR/ECAR between sexes (Fig. A-C). Notably, NDNs exhibited predominantly flat OCR traces in response to mitochondrial inhibitors, consistent with prior reports that neutrophils have minimal reliance on oxidative phosphorylation[25]. While maximal respiration was comparable, a trend suggested higher ATP production in male NDNs (p = 0.07, Fig.2C).

### Paired Analysis Reveals Subset-Specific Sexual Dimorphism

When comparing matched LDNs and NDNs within each sex, male LDNs had higher OCR than NDNs, where as for females the inverse was true, i.e. NDNs had higher OCR than LDNs (Fig. 3G and H). Male LDNs also showed higher glycolysis than their NDNs, whereas this difference was absent in females (Fig. 3 I and J).

## Discussion

Our study provides novel evidence of sex-specific metabolic programming in human neutrophil subsets and reveals that LDNs from males exhibit significantly higher oxidative metabolism than female LDNs, while maintaining comparable glycolytic activity. This divergence was subset-specific, as NDNs showed no sex difference in basal OCR but demonstrated a trend toward higher ATP production in males. Notably, paired analysis revealed a striking inversion in metabolic hierarchy between sexes where male LDNs displayed higher OCR than their matched NDNs, whereas female NDNs were more oxidative than their LDNs. These findings significantly extend previous work which described estrogen’s influence on neutrophil bioenergetics, by demonstrating that sexual dimorphism manifests differently across the total neutrophil population

The observed metabolic differences likely reflect fundamental sex hormone signaling pathways but also increased mitochondrial mass in male neutrophils, and agrees with previous results showing increased OCR in the total neutrophil population from young males[26]. This oxidative phenotype of male LDNs aligns with known androgen effects on mitochondrial biogenesis and efficiency [27], while the more glycolytic tendency of female LDNs may reflect estrogen-mediated promotion of aerobic glycolysis, similar to the Warburg effect observed in other immune cell types[28]. This metabolic stratification could have important functional consequences, as oxidative metabolism typically supports rapid antimicrobial responses [29], while glycolytic metabolism sustains prolonged inflammatory activity[30]. Our data suggest that even when neutrophil counts are equivalent between sexes, as we observed for both LDNs and NDNs, their functional metabolic states differ substantially.

Not only does this work expand our understanding of neutrophil biology, but these findings also carry important implications for understanding sex disparities in disease susceptibility and outcomes. LDNs play key roles in the pathogenesis of systemic lupus erythematosus [31], COVID-19 [32], breast cancer metastasis[18], and sepsis [33]. In female-predominant autoimmune conditions like lupus or rheumatoid arthritis, the glycolytic bias of female LDNs could potentiate persistent inflammation through enhanced NETosis or cytokine production. Conversely, in male-predominant conditions like severe COVID-19 or sepsis [34, 35], the heightened oxidative capacity of male LDNs may contribute to excessive tissue damage during hyperinflammatory responses. The subset-specific nature of these differences is particularly clinically relevant, as current diagnostic and therapeutic approaches typically treat neutrophils as a homogeneous population.

### Perspectives and Significance

From a therapeutic perspective, our work suggests that immunometabolic interventions targeting neutrophils may require sex-specific approaches. For instance, inhibitors of glycolysis might prove more effective in female-predominant inflammatory diseases, while mitochondrial-targeted therapies could be more relevant for male-predominant conditions. Several important considerations emerge from this work. The dynamic nature of neutrophil phenotypes suggests that metabolic states may shift during disease progression, warranting longitudinal studies. Finally, our findings underscore the necessity of sex-stratified analyses in both basic and clinical immunology research, as unisex approaches may obscure biologically and clinically relevant differences.

## Conclusions

Future studies should explore the mechanistic drivers of these metabolic differences, particularly the roles of androgen and estrogen receptor signaling in regulating neutrophil subset metabolism. Additionally, functional studies linking these metabolic profiles to specific immune activities like phagocytosis, NETosis, and cytokine production will be crucial for understanding their pathophysiological significance. Our work establishes a foundation for such investigations while highlighting the importance of considering both sex and neutrophil heterogeneity in immunological research and therapeutic development

## Declarations

### Ethics approval and consent to participate

Sample collection was approved by the St James’s Hospital/Tallaght University Hospital Joint Research Ethics Committee and all donors provided informed consent.

## Availability of data and materials

### Competing interests

The authors declare that they have no competing interests

### Funding

This work was funded by the Royal City of Dublin Hospital Trust and the Health Research Board (HRB-EIA-2024-002).

### Authors’ contributions

AY collected and analysed data and wrote the manuscript. FM collected data and wrote the manuscript. JK acquired funding and edited and reviewed the manuscript. GL acquired funding, designed the study, wrote and reviewed the manuscript.

## Acknowledgements

We would like to thank Dr. Lorraine Thong, Dr. Kevin Brown and Dr. Seán O’Donohue for their assistance in collecting samples from healthy donors for this manuscript.

## Notes

### Competing Interest Statement

The authors have declared no competing interest.

## References

1. Manuel, R.S.J. and Y. Liang, Sexual dimorphism in immunometabolism and autoimmunity: Impact on personalized medicine. Autoimmunity Reviews, 2021. 20(4): p. 102775.

2. Billi, A.C., J.M. Kahlenberg, and J.E. Gudjonsson, Sex bias in autoimmunity. Curr Opin Rheumatol, 2019. 31(1): p. 53–61.

3. Klein, S.L. and K.L. Flanagan, Sex differences in immune responses. Nature Reviews Immunology, 2016. 16(10): p. 626–638.

4. Conrad, N., et al., Incidence, prevalence, and co-occurrence of autoimmune disorders over time and by age, sex, and socioeconomic status: a population-based cohort study of 22 million individuals in the UK. The Lancet, 2023. 401(10391): p. 1878–1890.

5. Gay, L., et al., Sexual Dimorphism and Gender in Infectious Diseases. Frontiers in Immunology, 2021. Volume 12 - 2021.

6. Goble, F.C. and E.A. Konopka, SEX AS A FACTOR IN INFECTIOUS DISEASE. Transactions of the New York Academy of Sciences, 1973. 35(4 Series II): p. 325–346.

7. Lezama-Davila, C.M., et al., Sex-associated Susceptibility in Humans with Chiclero’s Ulcer: Resistance in Females is Associated with Increased Serum-levels of GM-CSF. Scandinavian Journal of Immunology, 2007. 65(2): p. 210–211.

8. !!! INVALID CITATION !!! [8-10].

9. Lee, J.H., J.P. Lydon, and C.H. Kim, Progesterone suppresses the mTOR pathway and promotes generation of induced regulatory T cells with increased stability. European Journal of Immunology, 2012. 42(10): p. 2683–2696.

10. Foo, Y.Z., et al., The effects of sex hormones on immune function: a meta-analysis. Biological Reviews, 2017. 92(1): p. 551–571.

11. Balogh, A., et al., Sex hormone-binding globulin provides a novel entry pathway for estradiol and influences subsequent signaling in lymphocytes via membrane receptor. Scientific reports, 2019. 9(1): p. 4.

12. Liang, Y., et al., A gene network regulated by the transcription factor VGLL3 as a promoter of sex-biased autoimmune diseases. Nature Immunology, 2017. 18(2): p. 152–160.

13. Hacbarth, E. and A. Kajdacsy-Balla, Low density neutrophils in patients with systemic lupus erythematosus, rheumatoid arthritis, and acute rheumatic fever. Arthritis & Rheumatism: Official Journal of the American College of Rheumatology, 1986. 29(11): p. 1334–1342.

14. Ssemaganda, A., et al., Characterization of neutrophil subsets in healthy human pregnancies. PloS one, 2014. 9(2): p. e85696.

15. Hardisty, G.R., et al., High purity isolation of low density neutrophils casts doubt on their exceptionality in health and disease. Frontiers in immunology, 2021. 12: p. 625922.

16. Tay, S.H., T. Celhar, and A.M. Fairhurst, Low-density neutrophils in systemic lupus erythematosus. Arthritis & Rheumatology, 2020. 72(10): p. 1587–1595.

17. Rosales, C., Neutrophil: A Cell with Many Roles in Inflammation or Several Cell Types? Frontiers in Physiology, 2018. Volume 9 - 2018.

18. Hsu, B.E., et al., Immature Low-Density Neutrophils Exhibit Metabolic Flexibility that Facilitates Breast Cancer Liver Metastasis. Cell Reports, 2019. 27(13): p. 3902-3915.e6.

19. Martin, K.R., et al., CD98 defines a metabolically flexible, proinflammatory subset of low-density neutrophils in systemic lupus erythematosus. Clin Transl Med, 2023. 13(1): p. e1150.

20. Ui Mhaonaigh, A., et al., Low Density Granulocytes in ANCA Vasculitis Are Heterogenous and Hypo-Responsive to Anti-Myeloperoxidase Antibodies. Front Immunol, 2019. 10: p. 2603.

21. Ishikawa, M., et al., Sex Differences of Neutrophil Extracellular Traps on Lipopolysaccharide-Stimulated Human Neutrophils. Surg Infect (Larchmt), 2024. 25(7): p. 505–512.

22. Svedlund Eriksson, E., et al., Testosterone exacerbates neutrophilia and cardiac injury in myocardial infarction via actions in bone marrow. Nature Communications, 2025. 16(1).

23. Blazkova, J., et al., Multicenter Systems Analysis of Human Blood Reveals Immature Neutrophils in Males and During Pregnancy. The Journal of Immunology, 2017. 198(6): p. 2479–2488.

24. Yennemadi, A.S., J. Keane, and G. Leisching, The Isolation and Characterization of Low- and Normal-Density Neutrophils from Whole Blood. JoVE, 2025(216): p. e67805.

25. Chacko, B.K., et al., Methods for defining distinct bioenergetic profiles in platelets, lymphocytes, monocytes, and neutrophils, and the oxidative burst from human blood. Laboratory investigation, 2013. 93(6): p. 690–700.

26. Gupta, S., et al., Sex differences in neutrophil biology modulate response to type I interferons and immunometabolism. Proceedings of the National Academy of Sciences, 2020. 117(28): p. 16481–16491.

27. Ahmad, I. and A.E. Newell-Fugate, Role of androgens and androgen receptor in control of mitochondrial function. American Journal of Physiology-Cell Physiology, 2022.

28. Cai, Q., et al., Regulation of glycolysis and the Warburg effect by estrogen-related receptors. Oncogene, 2013. 32(16): p. 2079–2086.

29. Kim, S.Y., et al., Antibacterial strategies inspired by the oxidative stress and response networks. Journal of Microbiology, 2019. 57(3): p. 203–212.

30. Soto-Heredero, G., Gómez de Las Heras MM, Gabandé-Rodríguez E, Oller J, Mittelbrunn M. Glycolysis-a key player in the inflammatory response. FEBS J, 2020. 287(16): p. 3350–69.

31. Rahman, S., et al., Low-density granulocytes activate T cells and demonstrate a non-suppressive role in systemic lupus erythematosus. Annals of the rheumatic diseases, 2019. 78(7): p. 957–966.

32. Morrissey, S.M., et al., A specific low-density neutrophil population correlates with hypercoagulation and disease severity in hospitalized COVID-19 patients. JCI insight, 2021. 6(9): p. e148435.

33. Sun, R., et al., Dysfunction of low-density neutrophils in peripheral circulation in patients with sepsis. Scientific reports, 2022. 12(1): p. 685.

34. Twitchell, D.K., et al., Examining male predominance of severe COVID-19 outcomes: a systematic review. Androgens: Clinical Research and Therapeutics, 2022. 3(1): p. 41–53.

35. Mester, P., et al., Exploring the relationship between plasma adiponectin, gender, and underlying diseases in severe illness. Biomedicines, 2023. 11(12): p. 3287.

